# Diversified agroforestry systems improve carbon foot printand farmer’s livelihood under limited irrigation conditions

**DOI:** 10.1101/2020.07.17.208405

**Authors:** Sanjay Singh Rathore, Kapila Shekhawat, VK Singh, Subhash Babu, RK Singh, PK Upadhyay, Ranjan Bhattacharyya

**Affiliations:** Division of Agronomy. ICAR-Indian Agricultural Research Institute (IARI), Pusa, New Delhi (India); Centre for Environment Science and Climate Resilient Agriculture ICAR-IARI, Pusa, New Delhi 110 012, India

**Keywords:** Agroforestry, diversification, system productivity, profitability, carbon sequestration

## Abstract

Increasing weather aberrations cause frequent crop failure in monoculture cropping system. Specialized crop production systems, where few seasonal crops occupy vast arable lands, resulting in more biotic and abiotic stresses in agri-ecosystem. Therefore a diversified agroforestry systemwas evaluated to ensure resilience underlimited water conditions, with an aim to augment carbon footprint with enhanced productivity and profitability. The study hypothesised that integration of perennial fruits trees with seasonal crops will have benign effect for sequestering more carbon and improving livelihood of the farmers. This is one of the first timesthat arid fruits tress along with leguminous,and other low water requiring crops were studied for improved carbon sequestration, livelihood of the farmers andfor better resilience in production system. The experimental findings showed that arid fruit trees along with leguminous, oilseeds and cash crops resulted in higher profitability and thus improved livelihood of the farmersin arid and semi-arid areas of South Asia. Diversified phalsa-mung bean-potato and moringa-mung bean-potato were the most productive agroforestry system (36.7t/ha and 36.2 t/ha respectively. Under limited irrigation conditions, Karonda (Carisa spp.)-mung bean potato system was found best in improving livelihood with maximum net return of $ 3529.1/ha with higher profitability/day ($ 19.9/day). Phalsa -MB-potato system was also recorded maximum water use efficiency (33.0 kg/ha-mm), whereas density of SOC was in Phalsa-cowpea-mustard (9.10 Mg/ha) and *moringa*-mung bean -potato AFS (9.16 Mg/ha). Carbon footprint analysis revealed that maximum net C gain was in Phalsa-mung bean -potato system (7030 Carbon equivalent kg CE/ha/year).

## Introduction

India is one of the leading nations in irrigated (68.38Mha) as well as in rainfed (86 Mha) arable lands globally (1). Escalating climatic risks are leading to huge loss to the farmers. Appropriate land-use systems which ensure resilience by minimizing the impact of climatic vulnerabilities arecrucial for livelihood security and are integral part of mitigation strategies to climate change. Agroforestry is one such land use system that may potentially support livelihood through simultaneous production of food, fodder and firewood as well as mitigation and adaptation to climate change. Due to changing climate scenarios, natural resource degradation is projected as a serious problem in the years to come (2). In this context, the perennial fruits trees have been identified usually more resilient to environmental constraints due to better capability to cope up with aberrant weather conditions and multifunctional biomass production even on marginal land (3) against annual crops.However,integration of seasonal crops with compatible fruit trees are key for successful cultivation in challenged ecosystem like arid and semi arid, also the performanceof different crop species varies depending on the growing conditions (4).Numerous studies revealed that fruit based agroforestrymodels (AHM) are the promising land use systems because these conservesoil and moisture, reduces soil erosion, sustains production and income at higher levels (4, 5,6). The system also increases carbon sequestration in the soil, due to decomposition of litter fall of fruit trees. This contributesin enhanced biological activity for the stability of rhizospheric environment. The agroforestrysystem is also supportive in generating additional employment, especially during off-season. It also provides scope for auxiliary industries like processing (preserves, jam, jelly, etc.), essential oils extraction, transportation, packaging, etc.Agroforestry systems have been identified potential solution for the twin climate and food security challenges (6).

Even in irrigated areas, due toincreasing biotic and abiotic stresses, ithas become a daunting task to achieve sustainably high farm productivity.Agroforestry systems involving trees and crops into fallow periods between two cropping seasonscan lead to higher crop yields in many parts of the tropics (7), and increased well-being of the farmers (8). This will certainly reduce the riskin farming even under stressed agri-ecosystems. Furthermore, with the adoption of location specific AFS as part of integrated farming system IFS) approaches will improve overall livelihood of the farmers. In India, 55% of arable lands are rainfed and this is the high time to make them sustainable, productive and risk free production systems. Single crop based management strategies will not help in achieving these goals under limited water ecologies, especially in rainfed areas. The integration of more agriculture and allied activities with field crops will certainly help in creating a sustainable agri-ecosystem.Integration of annual crops with fruit trees yields manifold benefits through secure production, income generation and restoration of ecosystem services (9-10) in a sustainable manner. Hence, AFS, including fruit based systems was evaluated under limited water conditions for food, nutrition and income and carbon neutral farming.

## Materials and methods

### Site characterization

The experiment was conducted at ICAR-Indian Agricultural Research Institute, New Delhi during 2015-18 (28.40° N, 77.12° E and 229 m elevation). The region has a semi-arid climate, with an average (of>30 years) annual rainfall of 650 mm (upto 80% of which received during July-September). Rainfall along the period of the cropping cycle (July to June) ranged from 533 to 1507 mm. The mean daily minimum temperature of 0–4°C in January, mean daily maximum temperature of 40–46°C in May-June and mean daily relative humidity of 67–83% during the experimentation years (Table1). The soil pH and EC were 7.8 and 0.32 dSm^-1^. The initial physical, chemical and biological properties of the soil is depicted in Table 1.Soil physico-chemical properties (soil texture, bulk density, infiltration rate, SOC, available N, available P and available K) were recorded to characterise soil site conditions. The texture of the soil was sandy loam with (44 %sand, 38 %silt and 18 % clay) with bulk density of 1.49-1.52 Mg/m^3^, whereas the infiltration ranged between 24.1-28.2 cm/hr. The soils were low in organic carbon content (0.30-0.34), medium in available P andK, while poor in available N (<250 kg/ha).

**Table 1.** Soil Physico-chemical characteristics of the experimental site

| AHS/Parameters | Soil texture | BD<br>Mg/m<sup>3</sup> | Infiltration,<br>mm/hr | SOC % | Available<br>N, kg/ha | Available<br>P<sub>2</sub>O<sub>5</sub>,<br>kg/ha | Available<br>K<sub>2</sub>O,<br>kg/ha |
| --- | --- | --- | --- | --- | --- | --- | --- |
| Karonda based AHS | Sandy loam | 1.51<sup>a</sup> | 26.5<sup>ab</sup> | 0.39<sup>ab</sup> | 152<sup>b</sup> | 18.5<sup>a</sup> | 295<sup>a</sup> |
| Phalsa based AHS | Sandy loam | 1.49<sup>a</sup> | 26.2<sup>ab</sup> | 0.40<sup>a</sup> | 149<sup>b</sup> | 17.6<sup>a</sup> | 298<sup>a</sup> |
| Moringa based AHS | Sandy loam | 1.50<sup>a</sup> | 28.2<sup>a</sup> | 0.42<sup>a</sup> | 170<sup>a</sup> | 16.9<sup>b</sup> | 301<sup>a</sup> |
| Guava based AHS | Sandy loam | 1.52<sup>a</sup> | 24.1<sup>b</sup> | 0.38<sup>b</sup> | 155<sup>b</sup> | 17.1<sup>ab</sup> | 299<sup>a</sup> |

Within a column, values represented with different lower-case letters indicate significant differences (P = 0.05).

### Experimental details

AFS was developed for round the year crop cultivation and generation of produce for regular income and employment. During kharif season, intercropping of legume crops were taken in the rows in between the fruit crops. In fruits crops phalsa (*Grewiaasiatica*), karonda (*Carissa carandas*), drum stick (*Moringaoleifera*), guava (*Psidiumguajava*) and pomegranate were grown. Among field crops during kharif season, vegetable cowpea, mung bean were cultivated as intercrops (Plate 1). The rain water is harvested and stored in the pond for life saving irrigation through micro irrigation system. The experimentation is on long term basis and data of four years have been pooled (2015-16 to 2018-19).

### Crop and fruits tree management

During rainy season, mung bean and cowpea were grown in the alleys of fruits tress. Samrat and kasha kanchan varieties of mungbean(MB) and cowpea (CP) were used, respectively. Samrat (MB) matures in 55-60 days, resistance to yellow mosaic virus (MYMV), whereas Kashi kanchanis (CP) ready for first harvest within 50-55 days. The sowing of both crops was done during the first week of July every year, cowpeas were harvested for vegetable pods and MB for seed yield. Seed rate and spacing were 10-12 kg/ha and 30×10 cm for both crops. The fertilizer N(25kg/ha), P_2_O_5_ (50 kg/ha) and K_2_O (40 kg/ha) were used. No irrigation was applied during kharif season. For plant protection against diseases and insect pest, standard practices were followed. During rabi (winter season) mustard (Pusa mustard 28) and potato (Kufari chipsona 1) were grown in alleys of fruit trees. The seed rate of mustard was 5 kg/ha with 30×10 cm spacing, whereas in case of potato 2500kg/ha tuberwere planted at 60×15 cm spacing.

### Litter fall and pruned biomass

Timely pruning of fruits tress were done for promotion of proper fruiting every year. Moringa, phalsa resulted in maximum biomass from pruning, which was done during relatively dormant phases (winter season) of each year. While in case of karonda and guava, because of lesser pruning requirement, comparatively less biomass (pruned) was harvested. Litter fall of all fruit tress was also collected annually with the help of litter traps of 1 × 1 m size placed at fourdirections under the tree during litter fall period of 10 months each year (January–December). Thus, in all 4 years litter fall was weighed using electronic balance after drying in oven and expressed in t/ ha.

**Plate 1.**
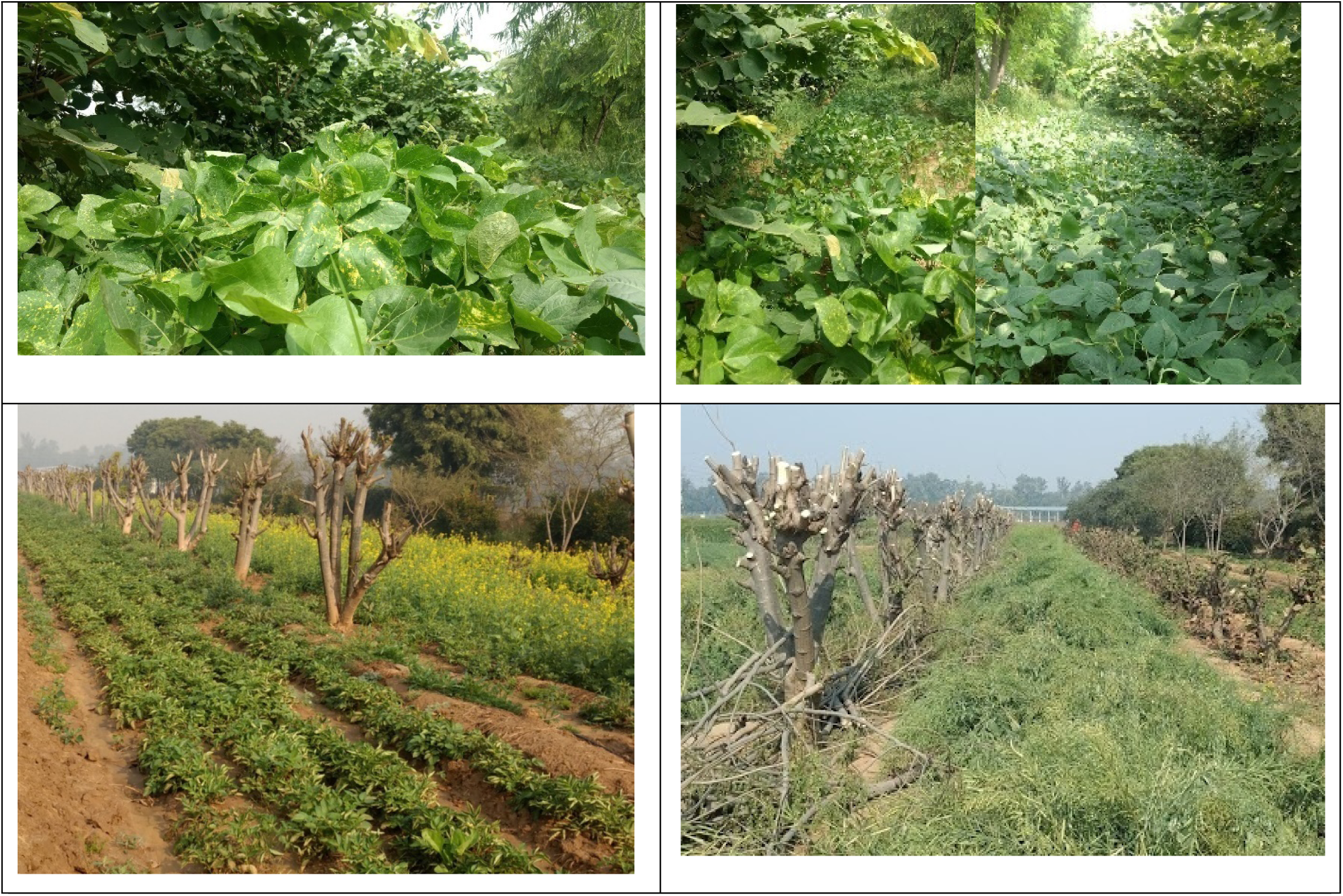
Mung bean, cowpea and mustard, potato as inter crops in Phalsa, Moringa, Karonda, aonla, guava based system for higher Carbon sequatration and enhanced system productvity and profitability

### Observations

Observations on growth, biomass, yield (seed, pod, fruits, wood) were recorded. The water usage was also estimated by quantifying the exact amount of irrigation and rainwater under different agroforestry systems. Carbon budgeting was also done by analysing change in carbon biomass and change in the soil carbon before start of the experiment and at the end of each harvest. Primary data on inputs (seeds, planting material, fertilizers, farmyard manure, pesticides, irrigation, and labour and machinery hours) utilized and outputs produced (crop, fruit yields and pruned wood) were recorded for each cropping season and annually of fruit plantations throughsystematic monitoring. The pruned biomass from each fruits tree was also measured separately. All inputs and outputs were converted into monetary values to present in acommon unit. For this, average price of each input/output over period was workedout to account for yearly price fluctuations. Governmentprices (minimum support price, MSP) were utilized if available. Mandi (local market) prices of commodities for which MSP were not available were used. Year-wise totals cost and total returns per hectare were pooled for each AFS to draw the inference of benefit: cost ratio. The economic return from the output was converted into $ based on the prevailing exchange rate of INR (Indian rupees) 75.5 for each $.

### Carbon sequestration potential (CSP)

CSP means an annual increment in the C sequestration by asystem under a particular treatment from a given base value. It wascalculated by the following expression:

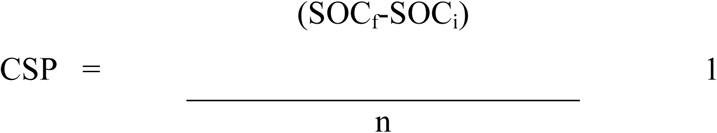

Where, SOC_f_ and SOC_i_ are the SOC (Mg/ha) in the final and initial soils, respectively, and n represents number of years of thestudy (11).

The C sequestration through different systems was calculated in terms of increases in C stock in soil. Data on initial and finalSOCconcentrations in the various treatments were collected for all plots.

Mass of soil organic carbon (MSOC) was estimated as below:

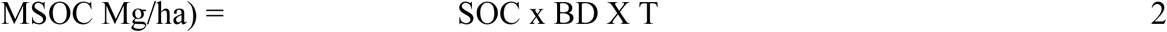

The MSOC in the soil layers (0–15, 15–30, and 30–45 cm) was calculated as Mg/ha where, MSOC is the mass of SOC (Mg/ ha), SOC is soil organic carbon concentration (%), BD is bulk density (Mg/ m^3^), and T is thickness of the surface layer (cm).

### Statistical analysis

All data generated on crop, fruits tress, economics, and other aspects were statistically analysed through SPSS and MS Excell and presented in different tables. Furthermore, the datawere analysed with descriptive statistics and one-wayANOVA at a significance level of 0.05. To computethe differences between the means, post hoc test wasperformed using Duncan’s Multiple Rangetest at significance level of0.05 by using SPSS (Statistical Package for SocialScience) version 20.0.

## Results

### Growth of annuals and perennials in AFS

Growth parameters of the intercrops (MB, CP, potato, mustard) were estimated, maximum AGR of MB (2.5 g day^-1^), potato (2.67 g day^-1^), CP (2.20 g day^-1^) and mustard (3.80 g day^-1^) was observed under moringa based AFS (Table 2). The AGR ranged between 1.8-3.6, 1.9-3.5, 2.2-3.8, 1.95-3.2 g day^-1^ in karonda, phalsa, moringa and guava based AFS, respectively.But almost all crops were recorded higher AGR under moringabased AFS, including CP. Among the field crops, maximum range in AGR was recorded in mustard (60 gday^-1^) and least was in case of MB (0.18 gday^-1^). Mustard had maximum mean AGR (3.83 g day^-1^). However, least AGR was noticed in CP due to its slow initial growth. Similar was the trend with CGR. Comparatively higher crop growth rate was observed in MB and among AFS,moringa basedAFShad higher CGR of different crops. The leaf area index (LAI) was higher in mustard (3.2-3.6) and least in potato (0.85-1.1).AFS of moringa and Phalsa resulted in higher LAI. As the leaves are the main photosynthetic parts in the plant, higher LAI indicate higher biomass accumulation and generally higher productivity. CGR varied from 6.83-12.3g/m^2^/day and it ranged between 11.5-13.1, 8.9-11.4, 10.5-12.8 and 6.2-7.9 g/m^2^/day for mungbean, potato, CP and mustard, respectively. Maximum variation was recorded in potato (2.5g/m^2^/day). The least mean CGR was recorded in mustard (6.83 g/m^2^/day) and maximum in MB (12.3 gm/m^2^/day). Leaves are the main photosynthetic organs in plants and LAI varied from 0.9-3.4. The mean LAI for MB, potato, cowpea and mustard was 1.68, 0.90, 1.74, 3.4, respectively. LAI varied more in cowpea (1.35-2.1) and variation in other field crops were comparatively lesser, but higher LAI was recorded in mustard crop (3.2-3.6).

**Table 2.**
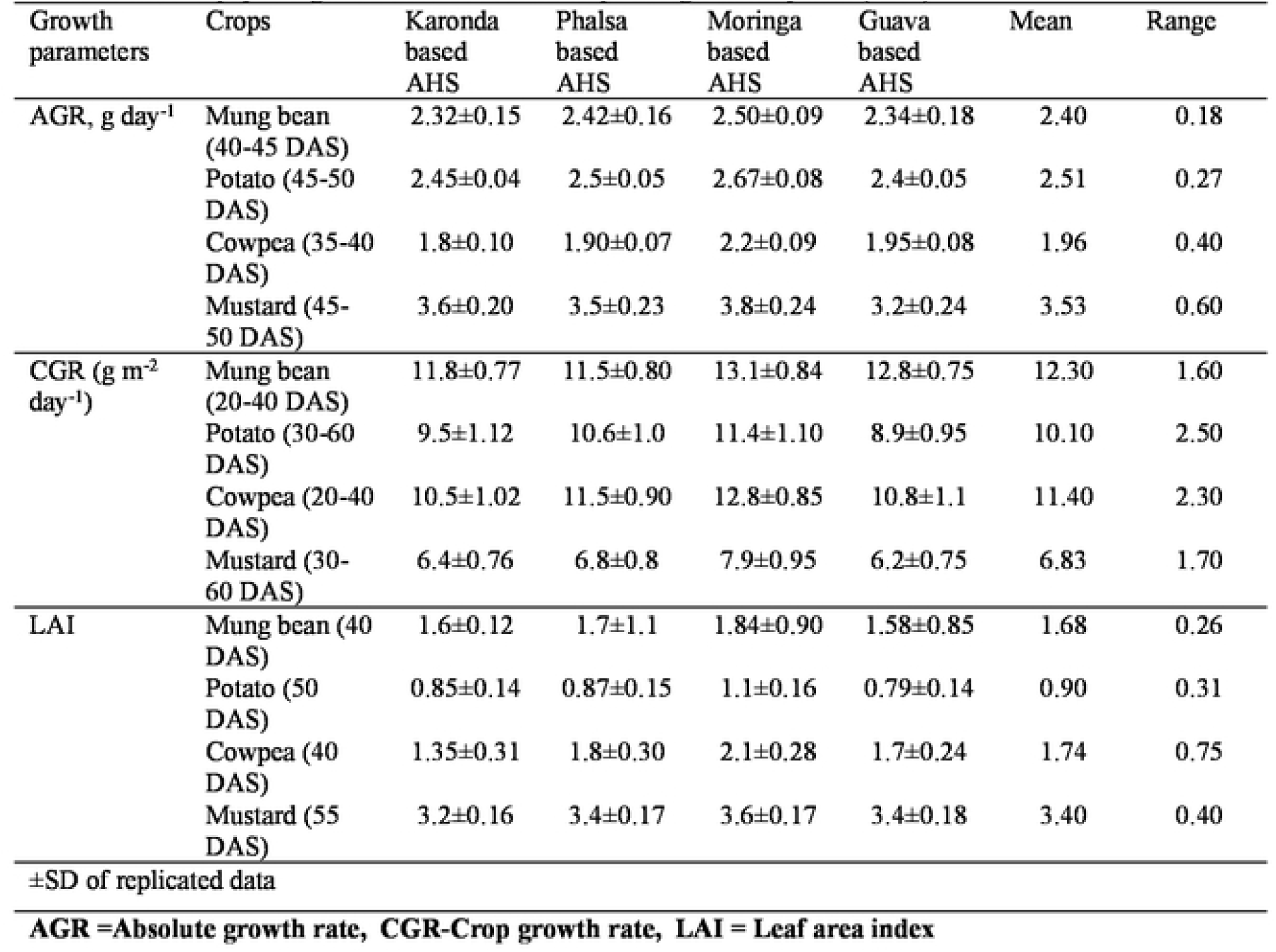
Growth/physiological attributes of field crops in Agri-horti system (AHS)

| Growth parameters | Crops | Karonda based AHS | Phalsa based AHS | Moringa based AHS | Guava based AHS | Mean | Range |
| --- | --- | --- | --- | --- | --- | --- | --- |
| AGR, g day<sup>-1</sup> | Mung bean (40-45 DAS) | 2.32±0.15 | 2.42±0.16 | 2.50±0.09 | 2.34±0.18 | 2.40 | 0.18 |
|  | Potato (45-50 DAS) | 2.45±0.04 | 2.5±0.05 | 2.67±0.08 | 2.4±0.05 | 2.51 | 0.27 |
|  | Cowpea (35-40 DAS) | 1.8±0.10 | 1.90±0.07 | 2.2±0.09 | 1.95±0.08 | 1.96 | 0.40 |
|  | Mustard (45-50 DAS) | 3.6±0.20 | 3.5±0.23 | 3.8±0.24 | 3.2±0.24 | 3.53 | 0.60 |
| CGR (g m<sup>-2</sup> day<sup>-1</sup>) | Mung bean (20-40 DAS) | 11.8±0.77 | 11.5±0.80 | 13.1±0.84 | 12.8±0.75 | 12.30 | 1.60 |
|  | Potato (30-60 DAS) | 9.5±1.12 | 10.6±1.0 | 11.4±1.10 | 8.9±0.95 | 10.10 | 2.50 |
|  | Cowpea (20-40 DAS) | 10.5±1.02 | 11.5±0.90 | 12.8±0.85 | 10.8±1.1 | 11.40 | 2.30 |
|  | Mustard (30-60 DAS) | 6.4±0.76 | 6.8±0.8 | 7.9±0.95 | 6.2±0.75 | 6.83 | 1.70 |
| LAI | Mung bean (40 DAS) | 1.6±0.12 | 1.7±1.1 | 1.84±0.90 | 1.58±0.85 | 1.68 | 0.26 |
|  | Potato (50 DAS) | 0.85±0.14 | 0.87±0.15 | 1.1±0.16 | 0.79±0.14 | 0.90 | 0.31 |
|  | Cowpea (40 DAS) | 1.35±0.31 | 1.8±0.30 | 2.1±0.28 | 1.7±0.24 | 1.74 | 0.75 |
|  | Mustard (55 DAS) | 3.2±0.16 | 3.4±0.17 | 3.6±0.17 | 3.4±0.18 | 3.40 | 0.40 |

±SD of replicated data
AGR =Absolute growth rate, CGR-Crop growth rate, LAI = Leaf area index

### Biomass and productivity

Integration of rainy season crops in AFS, resulted in higher green pod yield of CP from karonda based AFS, which was closely followed by *moringa*-CP-AFS (Table3). Among fruits trees, phalsa and moringa produced maximum fruits (13.0-14.5 t/ha) and least was from guava (Table 3). Seed yield of MB, a leguminous complementary crop in AFS, produced maximum underphalsa-based AFS. Moreover, mungbean under moringa-basedAFS was also resulted in almost equal seed productivity. The system productivities ofphalsa–CP (20.5 t/ha) and moringa-CP (19.6 t/ha) AFSwerehigherthan remaining combinations ofagroforestry systems during rainy seasons.Potato and mustard were taken as intercrops in AFS during rabi seasons. The tuber potato resulted in higher productivity ofM*oringa* based AFS (22.5 t/ha), closely followed by Karonda-potato system (22.0 t/ha). The seed yield of mustard was 1.6-1.75 t/ha (Table 3). Thus every drop of water is efficiently utilized for production of different crops. The data revealed thatphalsa-MB-potato was the most productive AFS (36.7t/ha), moringa-MB-potato was also have higher system productivity (36.2 t/ha). In these two AFS, fruit yieldswere also higher, which resulted in higher system productivity. Maximum system productivity was obtained from phalsa-mungbean-potato agroforestry system, which was closely followed by *moringa*-mungbean-potato system. The different AFS are economically viable, productive and employment generator round the year.

**Table 3.** Productivity of diversification with kharif, rabi crops in agri-horti system(pooled over three years)

| Fruits component | Fruit yield, t/ha | Kharif (rainy) season |  |  |  | Rabi (winter) season |  |  |  | System productivity and sustainability |  |  |
| --- | --- | --- | --- | --- | --- | --- | --- | --- | --- | --- | --- | --- |
|  |  | Seasonal crops | Green Pod yield (t/ha) | Seed Yield, t/ha | PE, kg/ha/day | Seasonal crops | Tuber yield (t/ha) | Seed yield, t/ha | PE, kg/ha/day | SP, t/ha | SPE, kg/ha/day | SYI |
| Karonda based | 5.0<sup>b</sup> | MB | - | 0.6<sup>b</sup> | 16.7<sup>d</sup> | Potato | 22 | - | 209.5a | 27.6<sup>b</sup> | 76<sup>c</sup> | 0.83<sup>a</sup> |
|  | 5.6<sup>b</sup> | CP | 7.5<sup>a</sup> | - | 35.6<sup>b</sup> | Mustard | - | 1.71 | 14.9b | 14.8<sup>d</sup> | 40<sup>d</sup> | 0.68<sup>ab</sup> |
| Phalsa based | 14.5<sup>a</sup> | MB | - | 0.8<sup>a</sup> | 41.7<sup>ab</sup> | Potato | 21.41 | - | 203.9a | 36.7<sup>a</sup> | 101<sup>a</sup> | 0.87<sup>a</sup> |
|  | 13.4<sup>a</sup> | CP | 7.2<sup>a</sup> | - | 56.3<sup>a</sup> | Mustard | - | 1.64 | 14.3b | 22.2<sup>bc</sup> | 61<sup>c</sup> | 0.79<sup>a</sup> |
| Moringa based | 13.0<sup>a</sup> | MB | - | 0.7<sup>ab</sup> | 37.5<sup>b</sup> | Potato | 22.5 | - | 214.3a | 36.2<sup>a</sup> | 99<sup>b</sup> | 0.87<sup>a</sup> |
|  | 12.4<sup>a</sup> | CP | 7.4<sup>a</sup> | - | 53.7<sup>a</sup> | Mustard | - | 1.71 | 14.9b | 21.5<sup>c</sup> | 59<sup>cd</sup> | 0.78<sup>a</sup> |
| Guava based | 2.2<sup>c</sup> | MB | 0.0 | 0.7<sup>ab</sup> | 6.8<sup>d</sup> | Potato | 21.1 | - | 201a | 24.0c | 66<sup>c</sup> | 0.80<sup>a</sup> |
|  | 1.6<sup>c</sup> | CP | 7.1<sup>a</sup> | - | 23.8<sup>c</sup> | Mustard | - | 1.75 | 15.2b | 10.4<sup>d</sup> | 29<sup>d</sup> | 0.55<sup>b</sup> |

Within a column, values represented with different lower-case letters indicate significant differences ( $P = 0.05$ ). MB: moong bean, CP: cowpea; SP = System productivity, t/ha, NR –Net returns, B:C ratio-Benefit: cost ratio Benefit :Cost ratio; PE = Production efficiency (kg/ha/day)

### Economics and profitability

With regards to economics, potato in AFS resulted in higher net returns ($1970.9-2156.1/ha). However, with inclusion of mustard in AFS, the net return was ($593.9-654.8/ha) higher than guava based AFS. The benefit: cost (B:C) ratio was >2.0 in potato based AFS in all fruit crops while remained lesser (1.87-2.06) in mustard integration with fruit trees (Table 4). However, lowest system productivity was in guava-CP-mustard AFS. With respect to profitability, maximum net system return was obtained from karonda-MB-potato system due to better prevailing market prices of the component commodities in the system. Similar was the trend in system profitability, system production efficiency. However, maximum B:C ratio was in karonda-MB-mustard system, as mustard involved comparatively lesser cost of cultivation. And highest cost of cultivation was with potato inclusion during rabi season. Overall under limited irrigation conditions, net return of $ 3529.1/ha with higher profitability/day ($ 9.67/ha/day) were observed in karonda-MB-potato system (Table 4).

**Table 4.**
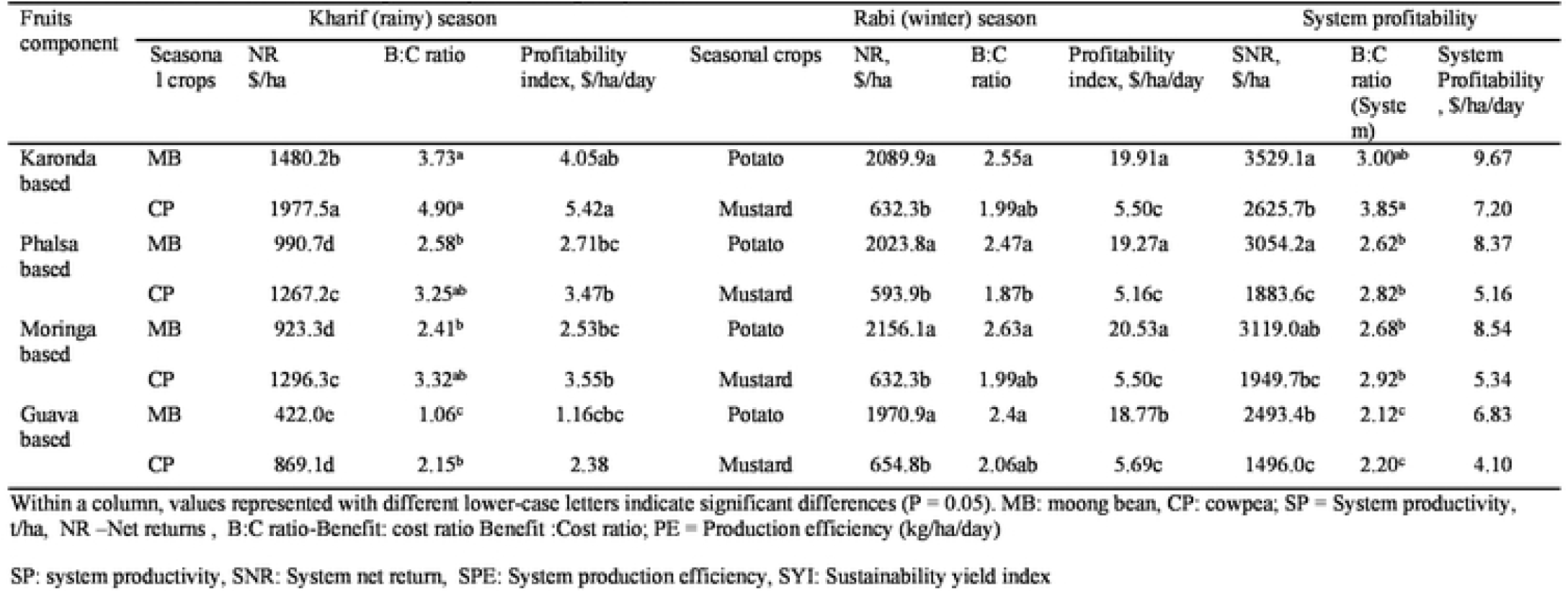
Economics of diversification in agri-horti system during rainy season

### Water use efficiency and water use dynamics

Water use under different AFS was 1112.4 mm and 1052.4 mm with respect to inclusion of potato and mustard during rabi (winter season), respectively in AFS (Table 5). System wise irrigation water was applied 180 mm in karonda, phalsa, moringa and guava based system with inclusion of potato, while 120 mm irrigation water was applied in fruit-MB/CP-mustard AFS. Kharif season crops (CP, MB) were grown as rainfed. Phalsa -MB-potato system recorded maximum WUE (33.0 kg/ha-mm) and IWUE (203.9 kg/ha-mm), closely followed by *Moringa*-MB-potato system (32.6 kg/ha-mm and 201.3 kg/ha-mm respectively).Contrary to WUE, different trend was observed in case of monetary irrigation water use efficiency (MIWUE) and karonda-CP-mustard systemresulted in maximumMIWUE (1654.2 kg/ha-mm), followed by karonda-MB-potato (1482.4 INR/ha-mm) and least was in guava-CP-mustard system (942.5 INR/ha-mm). Similar was the trend with MWUE (Table 5; Fig 1). Thus, phalsa-MB-potato agroforestry system was the most efficient system in terms of water use. However, monetary return per mm of water use was maximum in karonda-MB–potato system both in case of MWUE and MIWUE.

**Table5.** Water use, water use efficiency of agri-horti system under limited irrigation conditions.

| Fruit tree | Crop component |  | Water use , mm |  |  | WUE | IWUE | MWUE | MIWUE |
| --- | --- | --- | --- | --- | --- | --- | --- | --- | --- |
|  | Kharif | Rabi | Rainfall,<br>mm | Irrigation<br>water, mm | Total<br>water use,<br>mm | kg/ha-mm | kg/ha-mm | INR/ha-mm | INR/ha-mm |
| Karonda based | Mungbean | Potato | 932.4 | 180.0 | 1112.4 | 24.8<sup>b</sup> | 153.3<sup>ab</sup> | 239.9<sup>a</sup> | 1482.4<sup>b</sup> |
|  | Cowpea | Mustard | 932.4 | 120.0 | 1052.4 | 14.0<sup>b<sup>c</sup></sup> | 122.9<sup>b</sup> | 188.6<sup>abc</sup> | 1654.2<sup>a</sup> |
| Phalsa based | Mungbean | Potato | 932.4 | 180.0 | 1112.4 | 33.0<sup>a</sup> | 203.9<sup>a</sup> | 207.5<sup>b</sup> | 1282.6<sup>bc</sup> |
|  | Cowpea | Mustard | 932.4 | 120.0 | 1052.4 | 21.1<sup>ab</sup> | 185.3<sup>b</sup> | 135.3<sup>c</sup> | 1186.3<sup>c</sup> |
| Moringa based | Mungbean | Potato | 932.4 | 180.0 | 1112.4 | 32.6<sup>a</sup> | 201.3<sup>a</sup> | 212.0<sup>ab</sup> | 1309.9<sup>ab</sup> |
|  | Cowpea | Mustard | 932.4 | 120.0 | 1052.4 | 20.3<sup>ab</sup> | 178.0<sup>b</sup> | 140.1<sup>c</sup> | 1228.3<sup>bc</sup> |
| Guava based | Mungbean | Potato | 932.4 | 180.0 | 1112.4 | 21.6<sup>ab</sup> | 133.6<sup>bc</sup> | 169.5<sup>bc</sup> | 1047.3<sup>c</sup> |
|  | Cowpea | Mustard | 932.4 | 120.0 | 1052.4 | 9.9<sup>c</sup> | 86.8<sup>c</sup> | 107.5<sup>d</sup> | 942.5<sup>d</sup> |

Within a column, values represented with different lower-case letters indicate significant differences (P = 0.05).
WUE: water use efficacy, IWUE: irrigation water use efficiency, MWUE: monetary Water use efficiency, MIWUE: monetary irrigation water use efficiency

**Fig 1.**
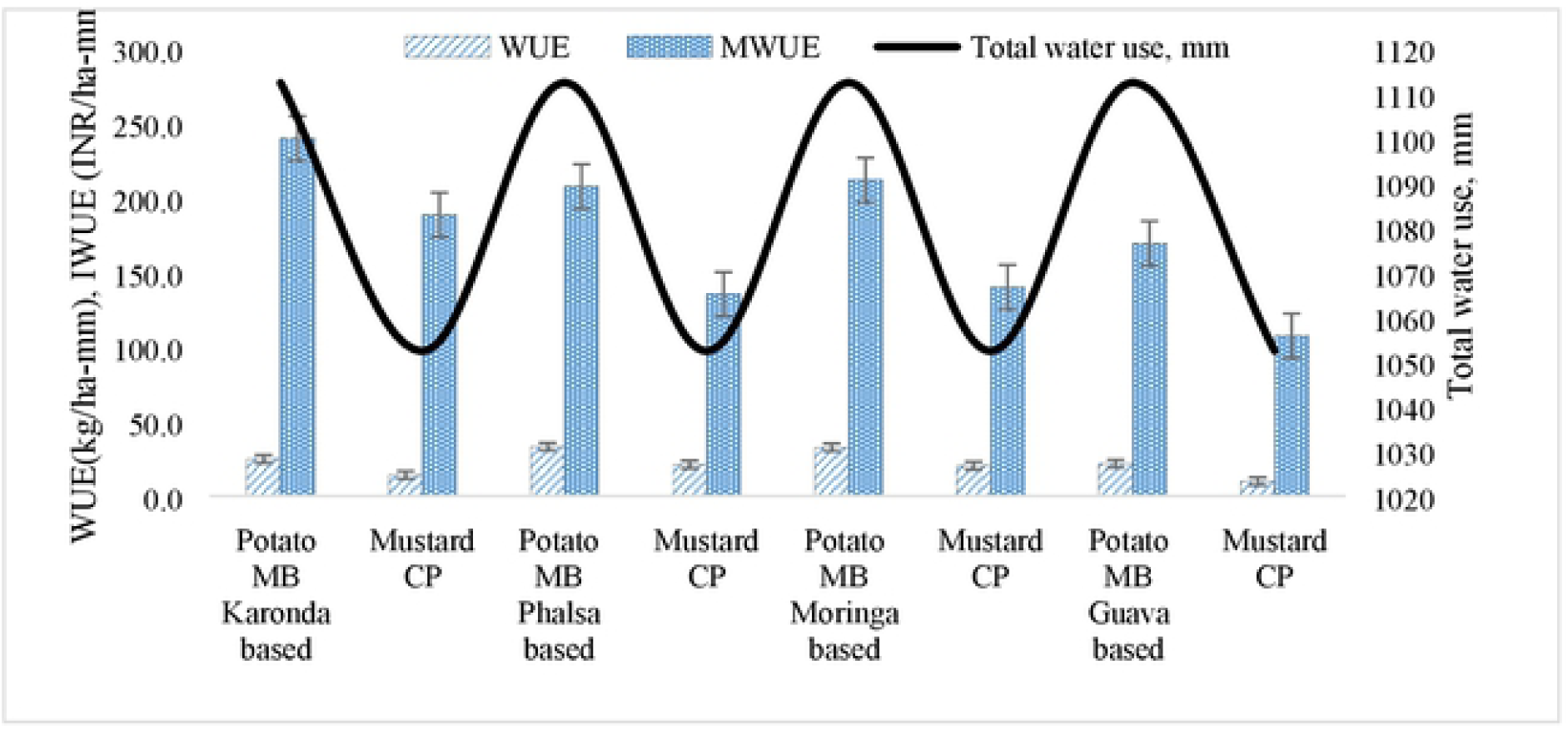
Water use efficiency (WUE), monetary water use efficiency (MWUE) and total water use under different agri-horti systems

### Carbon sequestration and carbon footprint

After third year of experimentation, maximum soil organic carbon(SOC) was in phalsa-CP-mustard (0.41%) and in *Moringa*-MB-potato system (0.41), even *Moringa*-CP-mustard was also recorded with similar SOC content (0.40%). However, least SOC was observed in guava-CP-mustard AFS (0.32) in the surface soil layers (0-15 cm). Slight variation was observed in the 15-30 cm of soil layer, karonda based AFS with MB-potato (0.33) and CP-mustard (0.32%), which was similar to guava based AFS (Table 6). Similar was the trend in mass of SOC and maximum density of SOC was in phalsa-CP-mustard (9.10 Mg/ha) and *moringa*-MB-potato AFS (9.16 Mg/ha). Carbon footprint analysis revealed that maximum net C gain was in phalsa-MB-potato system (Table 7) and least was in guava and karonda based AFS. After Phalsa based AFS, *Moringa* also resulted in significant higher build-up of carbon in soils. Among AFS, Phalsa based AFSresulted in maximum carbon sequestration rate potential (Fig 2).It is also evident that in surface soil (0-15 cm) higher carbon sequestration rate potential (CSRP) was observed (0.48-0.73 Mg/ha/year), whereas in deeper soils (15-30 cm) it ranged between 0.4-0.62 Mg/ha/year (Fig 3). Maximum CSRP was in Phalsa based AFS, which was due to higher biomass addition in soil from the system. Most of the organic residues remain on soil surface and it decreases with increase in soil depth due to lesser accumulation in the soil layers. Even in deeper soil layer (15-30 cm), CSRP was higher in Phalsa based AFS than it was in surface soil ofkaronda, *Moringa* and guava based AFS. The least sequestration of carbon was noticed in guava based AFS. But overall, *Moringa* based AFS was found to be the second best and among four AFSin terms of sequestration of carbon.

**Table6.** Soil organic carbon dynamics over the period (after third year of experimentation)

| Agrihorti system (AHS) |  |  | SOC (percentage) |  | MSOC (Mg/ha) |  | CSP (Mg/ha/yr) |  |
| --- | --- | --- | --- | --- | --- | --- | --- | --- |
| Fruits | Kharif crops | Rabi crops | 0-15 cm | 15-30 cm | 0-15 cm | 15-30 cm | 0-15 | 15-30 |
| Karonda based | Mungbean | Potato | 0.39<sup>ab</sup> | 0.33<sup>c</sup> | 8.72<sup>ab</sup> | 7.52<sup>b</sup> | 0.45<sup>c</sup> | 0.38<sup>c</sup> |
|  | Cowpea | Mustard | 0.39<sup>ab</sup> | 0.32<sup>c</sup> | 8.72<sup>ab</sup> | 7.34<sup>b</sup> | 0.52<sup>b</sup> | 0.38<sup>c</sup> |
| Phalsa based | Mungbean | Potato | 0.40<sup>ab</sup> | 0.36<sup>a</sup> | 8.72<sup>ab</sup> | 8.26<sup>a</sup> | 0.60<sup>ab</sup> | 0.76<sup>a</sup> |
|  | Cowpea | Mustard | 0.41<sup>a</sup> | 0.34<sup>b</sup> | 9.10<sup>a</sup> | 7.8<sup>o</sup> | 0.67<sup>a</sup> | 0.54<sup>b</sup> |
| Moringa based | Mungbean | Potato | 0.43<sup>a</sup> | 0.37<sup>a</sup> | 9.16<sup>a</sup> | 8.09<sup>a</sup> | 0.52<sup>b</sup> | 0.46<sup>b</sup> |
|  | Cowpea | Mustard | 0.42<sup>a</sup> | 0.36<sup>a</sup> | 8.94<sup>ab</sup> | 8.26<sup>a</sup> | 0.37<sup>c</sup> | 0.46<sup>b</sup> |
| Guava based | Mungbean | Potato | 0.39<sup>ab</sup> | 0.36<sup>ab</sup> | 8.1<sup>b</sup> | 7.34<sup>b</sup> | 0.44<sup>bc</sup> | 0.54<sup>b</sup> |
|  | Cowpea | Mustard | 0.37<sup>b</sup> | 0.35<sup>b</sup> | 7.6<sup>c</sup> | 6.89<sup>bc</sup> | 0.30<sup>d</sup> | 0.54<sup>b</sup> |

Within a column, values represented with different lower-case letters indicate significant differences (P = 0.05).

**Table 7.** Carbon foot print(Carbon equivalent kg CE/ha/year) in different agri-horti systems (AHS) under limited irrigation conditions.

| Agri-horti systems (AHS) |  |  | Fertilizer |  |  |  | Pesticides |  | Electricity | Fossil fuels | Total CE | Carbon output | Net C gain |
| --- | --- | --- | --- | --- | --- | --- | --- | --- | --- | --- | --- | --- | --- |
| Fruit tree | Crop component |  | N (1.3) | P (0.2) | K (0.15) | Total CEF | Insecticides | Fungicides |  |  |  |  |  |
| Karonda based | MB | Potato | 175.5<sup>b</sup> | 18<sup>c</sup> | 6.8<sup>b</sup> | 200.2<sup>b</sup> | 12.8<sup>b</sup> | 7.8<sup>a</sup> | 38.4 | 47.0 | 306.2<sup>ab</sup> | 2585<sup>c</sup> | 2279<sup>c</sup> |
|  | CP | Mustard | 175.5<sup>b</sup> | 18<sup>c</sup> | 6.8<sup>b</sup> | 200.2<sup>b</sup> | 12.8<sup>b</sup> | 3.9<sup>b</sup> | 28.8 | 35.2 | 280.9<sup>b</sup> | 2378<sup>c</sup> | 2097<sup>c</sup> |
| Phalsa based | MB | Potato | 188.5<sup>b</sup> | 18<sup>c</sup> | 6.8<sup>b</sup> | 213.2<sup>b</sup> | 15.3<sup>a</sup> | 7.8<sup>a</sup> | 38.4 | 47.0 | 321.8<sup>ab</sup> | 7352<sup>a</sup> | 7030<sup>a</sup> |
|  | CP | Mustard | 188.5<sup>b</sup> | 20<sup>bc</sup> | 6.8<sup>b</sup> | 215.2<sup>ab</sup> | 15.3<sup>b</sup> | 3.9<sup>b</sup> | 28.8 | 35.2 | 298.5<sup>b</sup> | 6660<sup>a</sup> | 6361<sup>a</sup> |
| Moringa based | MB | Potato | 84.5<sup>c</sup> | 22<sup>b</sup> | 7.5<sup>b</sup> | 114.0<sup>c</sup> | 12.8<sup>b</sup> | 7.8<sup>a</sup> | 38.4 | 47.0 | 220.0<sup>bc</sup> | 6232<sup>a</sup> | 6012<sup>a</sup> |
|  | CP | Mustard | 84.5<sup>c</sup> | 24<sup>ab</sup> | 7.5<sup>b</sup> | 116.0<sup>c</sup> | 12.7<sup>b</sup> | 3.9<sup>b</sup> | 28.8 | 35.2 | 196.7<sup>c</sup> | 5626<sup>b</sup> | 5429<sup>b</sup> |
| Guava based | MB | Potato | 227.5<sup>a</sup> | 26<sup>a</sup> | 12.0<sup>a</sup> | 265.5<sup>a</sup> | 15.3<sup>a</sup> | 7.8<sup>a</sup> | 38.4 | 47.0 | 374.0<sup>a</sup> | 2433<sup>c</sup> | 2059<sup>c</sup> |
|  | CP | Mustard | 227.5<sup>a</sup> | 28<sup>a</sup> | 12.0<sup>a</sup> | 267.5<sup>a</sup> | 15.3<sup>a</sup> | 3.9<sup>b</sup> | 28.8 | 35.2 | 350.7<sup>a</sup> | 2171<sup>c</sup> | 1821<sup>c</sup> |

Within a column, values represented with different lower-case letters indicate significant differences (P = 0.05).

**Fig 2.**
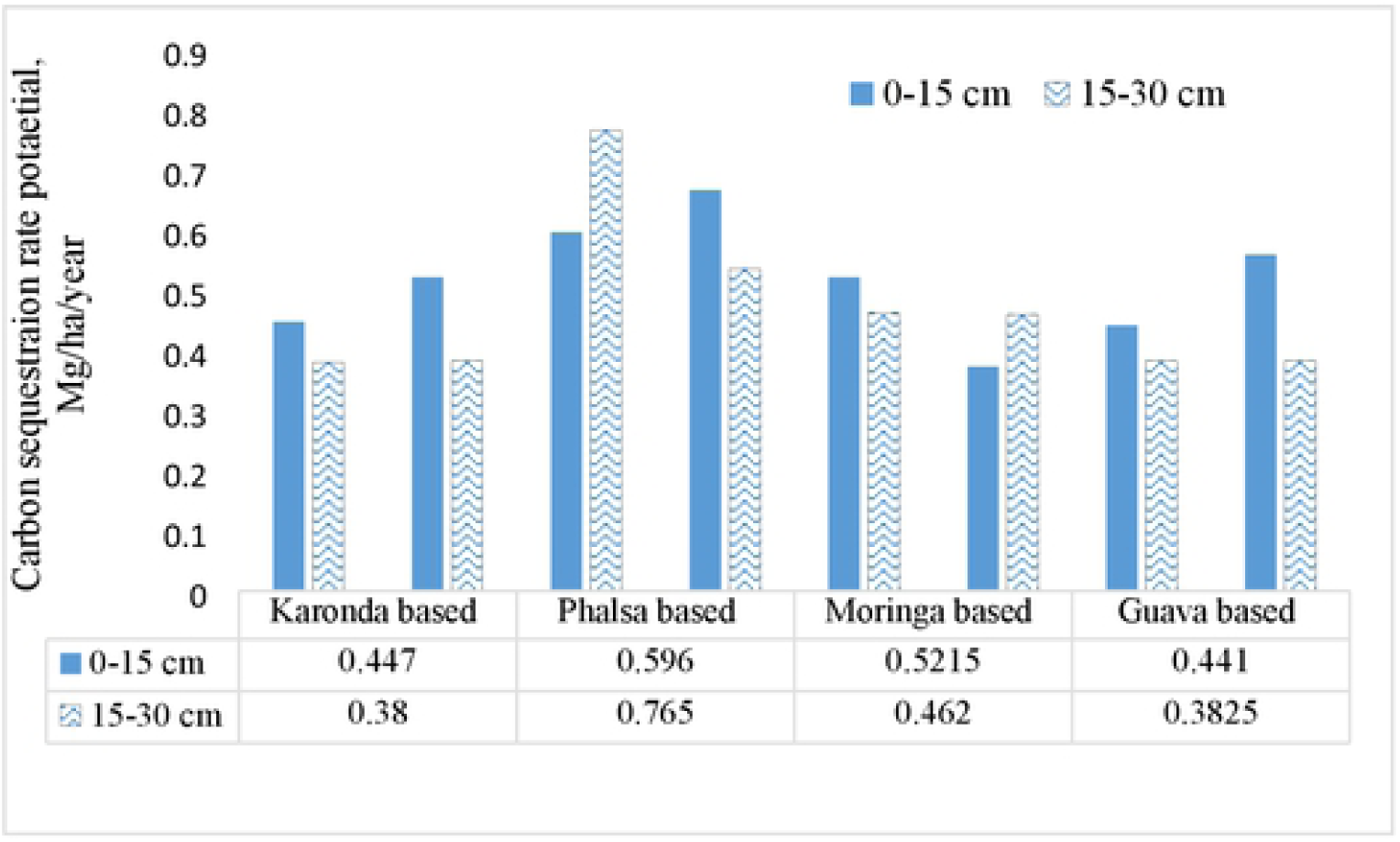
Carbon sequestration rate potential, Mg/ha/yr (CSRP) under fruit based system

**Fig 3.**
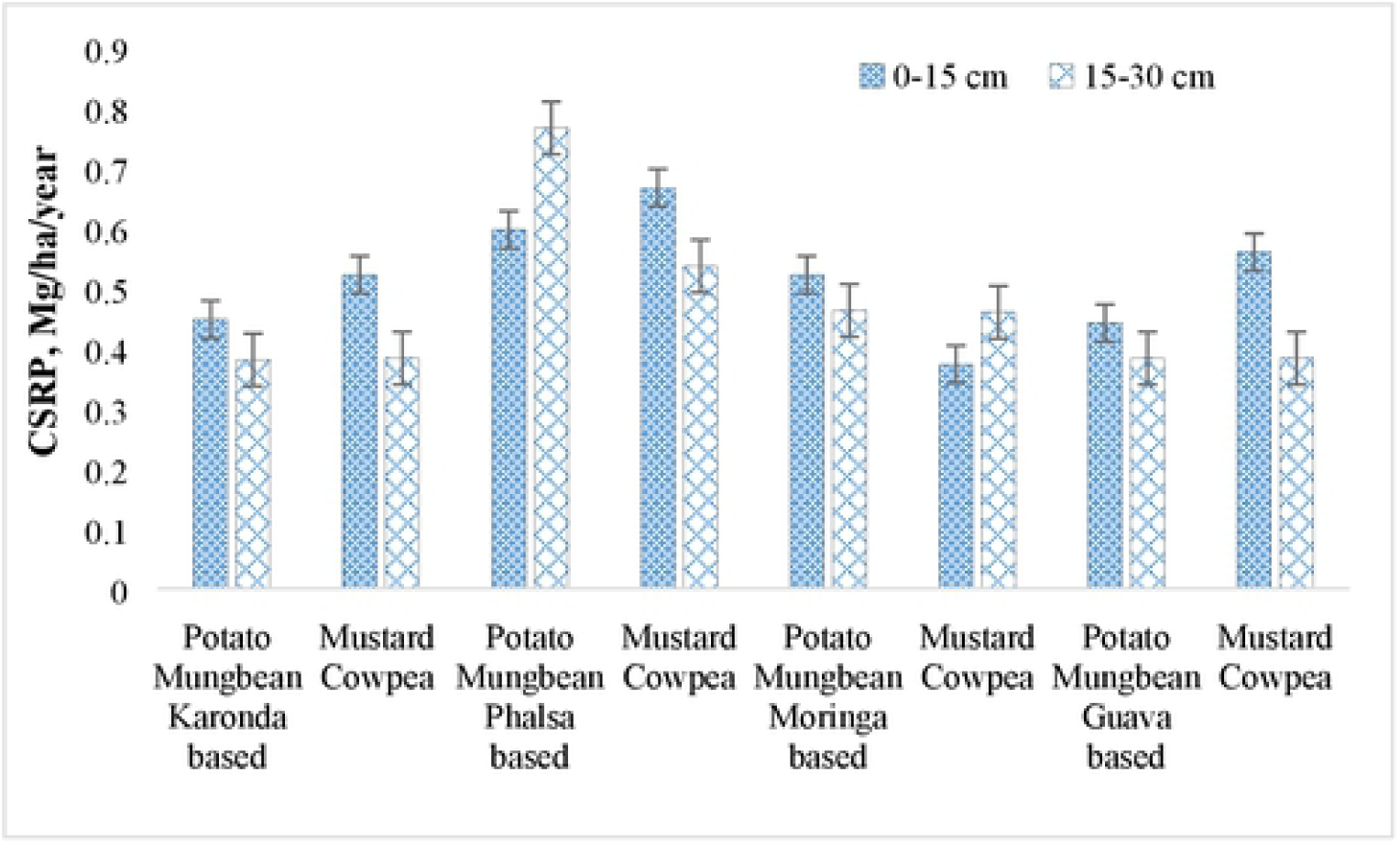
Carbon sequestration rate potential (CSRP) of agri horti systems

## Discussion

### Growth of annuals and perennials in AFS

The growth parameters of the field crops were found higher in *Moringa* based AFS. This was due to the fact that *Moringa*is multi propose trees, it fixes substantial amount of atm –N in to the soil (14 -15 13). There was less competition between the trees and the field crops, this ultimately resulted in higher biomass accumulation and better canopy spread in terms of higher LAI (Table 2). Maximum AGR of MB (2.5 g day^-1^), potato (2.67 g day^-1^), cowpea (2.20 g day^-1^) and mustard (3.80 g day^-1^) was observed under MoringabasedAFS (Table 2). Similarly, comparatively higher crop growth rate was observed in MB and among agri-horti systems (AFSs), *Moringa* based AFS recorded higher CGR of different crops. Foid et al. (14) spelled out the significance of *Moringa* in tropical and subtropical world, as it is a multipurpose tree, characterized by high biomass yield, and can tolerate unfavourable environmental conditions. The leaf fall and pruned biomass fall contributed immensely in building soil fertility which was reflected in growth and other productivity traits of different crops. *Moringa*trees are also valuable for alley cropping systems because they fix nitrogen which is supplied to intercrops via the pods and leaves they produce (15).Among the field crops, maximum range in AGR was recorded in mustard (0.60 gday^-1^) and least was in case of MB (0.18 gday^-1^). The effect of shedding under AFS on mustardvaried due to the fact of differential response of plant to rhizospheric and aerial environment, and also some tree species led to allelopathic effect on mustard which suppress plant growth (16,11). The leaf area index (LAI) was higher in case of the mustard crop (3.2-3.6) and least was in the potato crop (0.85-1.1), AFS of *Moringa* and phalsa resulted in higher LAI. As the leaves are the main photosynthetic parts in the plant, higher LAI indicate higher biomass accumulation and generally higher productivity. LAI varied from 0.9-3.4. Some tress like Guava have allelapathic effect which often result in lower crop growth, leaf area, LAI in agroforestry systems (19).

### Biomass and productivity

MB and CP intercropping in karonda basedAFS produced higher crop as well fruit yield, which was closely followed by *Moringa*-CP-AFS (Table3). *Moringa* tree, help in building soil fertility but its canopy spread was more than karonada and its shedding effect hinders adequate plat growth (13,17). This might be due to less canopy spread of karonda and least shedding effect on the intercrops.Also tree canopy of *Moringa* has an umbrellashaped crown with bi-(tri-) pinnate leaves, while the individual leaflets have a leaf area of one to two cm^2^, but higher canopy area (18). Singh et al. (**19**) also explained adaptive advantages of multipurpose karonda, rich in iron, widely grown as live fencing under arid and semi-arid regions, a hardy species and can tolerate extreme weather conditions.Karonda is an evergreen spiny shrub or a small tree up to 3 m height and suitable for arid tropics and sub-tropicshaving less requirement of inputs, grows successfully on marginal lands. It yields a heavy crop of attractive berry like fruits which are edible and rich in vitamin C and minerals especially iron, calcium, magnesium and phosphorus and good foragroforestry system (20). MB another leguminous intercropwasgrown in AFS produced maximum under phasla based system. Maximum system productivity was obtained from Phalsa-MB-potato and *Moringa*-MB-potato AFS. Pareek and Awasthi(**21**) spelled out higher system productivity under Phalsa based AFS and higher intercrop yield due to less competition for nutrients and water and recommended the synergy of phalsa in AFS. This was due to the fact that more space was available in alleys of Phalsa, causing no shedding effect on crops. Also good amount of leaf fall was added in the system (22). Hence,agroforestry systems are economically viable, productive and also create employment round the year.

### Economics and profitability

Inclusion of cash crop like potato in AFS resulted in higher net returns compared to mustard in AFS. But comparatively lesser cost of cultivation was involved in mustard over potato system (23).Hence the investment capacity of the farmersalso decide the system net returns. Both crops have benign relationship and this led to highersystem productivity in AFS. The B:Cratio was >2.0 in potato based AFS in all fruit crops, while it remained lesser in mustard integration with fruit trees. 26-27 Rathore and Bhatt (24) and Rathore et al. (25) also reported integration of suitable tree species with intercrops of complementary in association, ensure higher profitability from a system. The lowest system productivity in Guava-CP-mustard AFSwas due to poor productivity of Guava orchard. With respect to profitability, maximum net system return was obtained from karonda-MB-potato system due to better prevailing market prices of the component commodities in the system. Similar was the trend in system profitability, system production efficiency. The maximum B: C ratio was in karonda-MB-mustard system. This was due to the fact that mustard involved comparatively lesser cost of cultivation (19, 23). And highest cost of cultivation was with potato inclusion during rabi season. Nankar(26), Lyngbaek et al. (2001) and Arnold (28) also reported higher cost of cultivation of potato, even under intercropping, leading to lower B:C ratio.

### Water use efficiency and water use dynamics

Water use was higher in AFS including potato crop, as the water requirement of potato is more (180 mm), while in case of mustard 120 mm water was sufficient (23). A 120 day potato crop (kufari chipsona) consumes from 400-500 mm of water, and a substantial amount of water is also contributed by soil profile during the crop growth period. As the crop is grown under water limiting conditions, hence only 1800 mm water as supplementing irrigation was applied in the system, however depletion of more than 50 percent of the total available soil water during the growing period results in lower yields (29). Phalsa -MB-potato system recorded maximum WUE (33.0 kg/ha-mm) and IWUE (203.9 kg/ha-mm), closely followed by *Moringa*-MB-potato system (32.6 kg/ha-mm and 201.3 kg/ha-mm,respectively). This was due to the fact that better use of water by the crops in terms of higher plant growth which also resulted in higher productivity. Contrary to WUE, different trendswere observed in case of monetary irrigation water use efficiency (MIWUE) and karonda-cowpea-mustard system resulted in maximum MIWUE (1654.2 kg/ha-mm), followed by karonda-MB-potato (1482.4 INR/ha-mm) and least was in guava-cowpea-mustard system (942.5 INR/ha-mm). Similar was the trend with MWUE (Table 4 and Fig 1). Under arid and semi-arid conditions,Rathore et al. (30) also reported higher WUE ofmustard crop. This was due to higher productivity of karonda based AFS and better prevailing market prices of the karonda fruits. Karonda is also superiorly adopted under limited irrigation conditions to other AFS. Sharma et al. (31), Singh et al. (19) also reported higher productivity and profitability and water use from karonda based AFS under arid conditions. Thus, Phalsa-MB-potato AFS was the most efficient system in terms of water use. However, monetary return per mm of water use was maximum in karonda-MB –potato system, both in case of MWUE and MIWUE. Therefore, it can be concluded that AFS including fruit tree karonda and *Moringa* with MB and potato can improve overall farm productivity and also the profitability under limited irrigation conditions.

### Carbon foot print and carbon sequestration

Soil organic carbon is one the key indicators for optimum soil and environmental sustainability (31-32 32-33), Nevertheless, Agroforestry has been recognized as having the greatest potential for C sequestration of all the land uses as significant amount of C stored in aboveground biomass, agroforestry systems can also store C belowground (34). Agroforestry system including perennial trees can increase carbon sequestration, offsetGHG emissions and reduce the carbon footprint generated by animal production (35). It was also evident from phalsa-CP-mustard (0.41%) and in *moringa*-MB-potato system (0.41), where maximum soil organic carbon (SOC) was noticed. Similarly, highermass of SOC was in phalsa-CP-mustard (9.10 Mg/ha) and *Moringa*-MB-potato AFS (9.16 Mg/ha). Among AFS, phalsa based AFS resulted in maximum carbon sequestration rate potential (Fig 2), especially in surface soil (0-15 cm). However, in deeper soils (15-30 cm) it ranged between 0.4-0.62 Mg/ha/year. Greater biomass addition in soil from the phalsa based AFS resulted in increased CSRP. Most of the organic residues remain on soil surface and it decreased with increase in soil depth due to lesser accumulation in these soil layers. Leaf fall as well pruned biomass was comparatively higher in Phalsa based system. Moring*a* based AFS has more nutrient especially leaf N but total biomass was lesser and it leads to reduced CSRP over phalsa based AFS. The noticeable role of AFS in carbon sequestration has increased global interest to stabilise GHG emissions (34 Rizvi et al., 2019). Furthermore, Diana et al. (36) reported that on average, carbon benefits are greater in agroforestry systems.Udawatta and Jose (37), 2011 precisely explained that in tree based systems, C sequestration could be enhanced in a short period of time due to enhanced soil C sequestration potential in alley cropping than in monocropping systems. This was due to the fact that lesser biomass accumulation from fruits and also from seasonal crops was observed in the system. Carbon footprint analysis revealed that maximum net C gain was in phalsa-MB-potato system (Table 6), and least was in guava and karonda based AFS. This was due to the fact that no pruned biomass (wood, etc.) was produced in karonda based AFS. After phalsa based AFS, *moringa* also resulted in significant higher gain of carbon (38-40).

## Conclusion

The study confirms that integration of perennial fruits tress with field crops (MB, CP, mustard and potato) have the potential to enhance system productivity (fruit, vegetables, seed), improve soil health with better carbon sequestration and lesser C footprint. Phalsa-MB-potato and *moringa*-MB-potato AFS resulted in maximum system productivity. Inclusion of cash crop like potato in AFS resulted in higher net returns than mustard in AFS. The B:C ratio was >2.0 in potato based AFS in all fruit crops, while remained lesser in mustard integration with fruit trees. With respect to profitability, maximum net system return was obtained from karonda-MB-potato system. Phalsa -MB-potato system recorded maximum WUE (33.0 kg/ha-mm) and IWUE (203.9 kg/ha-mm), due to better use of water by the crops in terms of higher plant growth which also resulted in higher productivity. However, monetary return per mm of water use was maximum in karonda-MB –potato system both in terms of MWUE and MIWUE. Phalsa-CP-mustard and in *Moringa*-MB-potato system were effective mitigation strategies against the emission of GHG. Phalsa-CP-mustard (0.41%) and in *Moringa*-MB-potato systemshad maximum soil organic carbon (>0.40%) and higher density of SOC (9.16 Mg/ha). Phalsa based AFS also resulted in maximum carbon sequestration rate potential, especially in surface soil (0-15 cm). The noticeable role of AFS in carbon sequestration has increased global interest to stabilise greenhouse gas (GHG) emissions.Carbon footprint analysis revealed that maximum net C gain was in phalsa-MB-potato system. Therefore, it can be concluded that that agroforestry system including fruit tree karonda and *Moringa* with MB and potato can improve overall farm productivity and also the profitability under limited irrigation conditions.

## Acknowledgement

The authors are highly grateful to the Director, ICAR-Indian Agricultural Research Institute (IARI), Head, Division of Agronomy for providing all the logistics to conduct this experiments. The contribution of technical and field staff is also dully acknowledged for successful conduct and timely record of all observations in the experiment.

